# Biological Organization, Biological Information, and Knowledge

**DOI:** 10.1101/012617

**Authors:** Maurício Vieira Kritz

## Abstract

A concept of information designed to handle information conveyed by organizations is introduced. This concept of information may be used at all biological scales: from molecular and intracellular to multi-cellular organisms and human beings, and further on to collectivities, societies and culture. In this short account, two ground concepts necessary for developing the definition will also be introduced: whole-part graphs, a model for biological organization, and synexions, their immersion into space-time. This definition of information formalizes perception, observers and interpretation; allowing for considering information-exchange as a basic form of biological interaction. Some of its elements will be clarified by arguing and explaining why the immersion of whole-part graphs in (the physical) space-time is needed.

## 1 Introduction

Organization is a key characteristic of biological entities and phenomena [11, 12]. It appears everywhere: from simple oscillatory chemical reactions and the structure of macro-molecules to cells’ activation-inhibition and consensus bio-chemical setups, moving molecular aggregates, modules and motifs with associated biological functions, and stable organelles. It expands beyond cell organelles and cell inner structures into tissues and organs of multicellular organisms and further upward into populations, societies and cultures, although its instantiation at each scale may present seemingly uncorrelated forms [26, 20].

Organization is also tightly intertwined with information. Any organization may convey some information to someone. Or else, an organization may mean something to someone. Moreover, different organizations like two words in different languages may convey the same information, while the same organization (like homonyms in spoken language) may possess distinct meanings. Analogous situations do occur at the sub-cellular level where and when the same molecular aggregates perform different biological functions. How can this be formalized? To what extend can formalized versions of these concepts clarify and identify the various relations between organization and information.

The purpose of this writing is to present a definition of (biologic) information based on organizations, to clarify some characteristics of organizations and to discuss the importance of considering space-time organizations in the definition of information. Hence, a definition of organization, a mathematical model for it, and organizations in space-time are presented in the next section. In the third section, the idea of perception in a very specific sense is introduced and the definition of information presented. In the fourth section, arguments and examples highlighting the importance of considering time and space in organizations to properly define biological information are presented at various scales and within different domains of discourse. The last section contains some concluding remarks and acknowledgements.

## 2 Biological Organization: a minimalist snapshot

This section provides a review of published material [15, 16] as well as new material. The former is included here for the sake of completeness and ready accessibility. The presentation in this section shortens technical details in favor of examples and clarifying explanations. The arguments are less formal and more illustrative, looking forward to throw light on the basal concepts, letting the reader more at easy with them.

### 2.1 Organizations

Consider a large enough heap of bricks indistinguishable with respect to all relevant attributes, characteristics and factors. The bricks are thus identical although not the *same*. This heap constitutes a set of bricks. Choose now 20 of these bricks at random. These 20 chosen bricks are still a set of bricks. They form a subset of the heap, if you don’t take the chosen bricks apart from the heap. If you do, there will be two heaps and their connection may become imperceptible.

Ensuite, pick all bricks in the heap and build a house. The house has 5 pieces: a living room, two bedrooms, a bathroom and a kitchen. From the living room there is a hall giving access to the bedrooms, the kitchen and the bathroom. Walls, made with the bricks, divide and delimit each piece. Some walls have doors, some other have windows, a few may have both. After the house is built, there are no more bricks but walls and rooms. Is a house a set of bricks? Is a brick in a wall identical to a brick in another wall or to another brick somewhere else in the same wall?

Consider any of the walls of the house. Mark 20 bricks in it at random. Are these bricks as indistinguishable as before? They may be side by side and taking them out will make a hole in the wall; luckily the hole may become a window or a door, depending on its height and shape. They may also be scattered throughout the wall; what may be a very refreshing strategy. Or they may concentrate in a junction of two walls, weakening its structure and ruining the house. The bricks are in consequence not indistinguishable anymore. They are certainly interchangeable (any two of them may be swapped…) but not indistinguishable; the effect of taking any (or a collection) of them out of the walls is not anymore the same. They became parts of a wall, which in turn became a part of the pieces of the house, and these latter parts of the house.

Organization is what distinguishes a house from a set of bricks. It may be informally defined in the following way. An atom here refers to an epistemological or cognitive atom — something we cannot, or do not want to, subdivide.

#### Definition 2.1

*An **Organization** is either:*

▪ *an atom;*
▪ *a set of organizations;*
▪ *a group of organizations put somehow in relation to one another;*
▪ *nothing else.*

This definition purposefully lacks a better clarification of the expression *somehow in relation with one another*, since this may be interpreted in several ways. Notwithstanding, it already include sets, of either atoms, organizations or both, as organizations.

In the case of the above example, for instance, bricks are atoms. Even if one is taken apart in the building process, we do not want to describe or analyze of what and how they are made off, independently of the relevance of their attributes. Nevertheless, we may consider them associated in many ways. A brick may be associated to another if they are in contact, sharing a face or a portion of a face, if they belong to the same wall, or yet to a specified region of a wall.

This idea of organization may be applied even to the heap itself. A brick may be associated to another if they are close together and groups of bricks may be considered separately. The ones at the top of the heap or at its base, for instance. In this case, however, shaking the heap a little could provoke a complete upheaval in this organization. In terms of organization, the heap is not as stable as a house, although the heap is pretty stable as a set.

### 2.2 Definition and Model

To make this definition more precise, establishing along a mathematical model for organizations, we will need to restraint what shall be considered as atoms in each situation to be modeled. All atoms of an organization shall be elements of an universal *admissible* set U. Using admissible sets allow us to consider certain sets as atoms without messing the organization construct [5, 13]. These sets differ in concept and formalism from the set in clause 2 of Definition 2.1.

This set U need not to be the same in different modeling situations, unless one want to compare the modeled organizations or make them interact somehow. It may also be typed; that is, U may be partitioned and its elements distributed among its partition classes. There is, in principle, no restriction on the number of elements of U or its type-classes. For instance, U can be a set of letters or letter groups while studying the organization of texts and of chemical atoms or substrates while considering the organization of molecules or biological entities. One may consider also an U consisting of both letters and chemical atoms if this helps representing the phenomena in hand.

Besides U, we will need some meta-elements: an enumerable set of meta-variables V = {v_0_, v_1_, v_3_,…} and a collection of special *undistinguished* elements ⊙. Meta-variables may be considered as voids in organizations — places where organizations may grow or refine — whatever appropriate. Elements ⊙ are generic nicknames for organizations that may become part of another, like in clauses 2 and 3 of Definition 2.1. Being indistinguishable allow for swapping equivalent (or indistinguishable) sub-organizations without changing the overall organization. Like the bricks in a wall.

Associations will be represented by hyper-graphs [7, 28]. However, to get things right, we need that its nodes be taken from the extended admissible set U_⊙_ = U ∪ V ∪ {⊙}. Hyper-graph with nodes from U_⊙_ will be called *extended hyper-graphs* to make clear that its nodes may be meta-elements. Whole-part graphs are constructed by binding extended hyper-graphs by “assigning” or “attributing” the generic ⊙ element of a hyper-graph *h*^*p*^ to a meta-variable in another hyper-graph *h*^*w*^. This construction start from the following operator prototype:

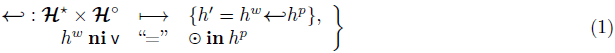

where 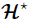 denotes the class of all extended hyper-graphs that have meta-variables v as nodes, 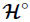 denotes the class of all extended hyper-graphs that have the special meta-element ⊙ as a node, and **ni** is the mirror of *u* **in** *h* that is a predicate deciding whether *u* is or is not a node of *h*.

An extended hyper-graph has just one ⊙ element as node (one nickname) but may have many meta-variables v_*i*_ as nodes. Moreover, if several hyper-graphs are to be bound to *h*^*w*^, there is no special reason to privilege any meta-variable to be bound before the others. Thus, the binding operator should be seen as a collateral binding of hyper-graphs 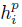 to *h*^*w*^:

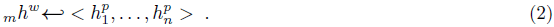

where *m* is the number of meta-variables in *h*^*w*^, *n* ≤ *m* is the number of extended hyper-graphs to be bound to *h*^*w*^. Without loss of generality, it is supposed that 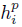 binds to v_*i*_.

Using the representation of hyper-graphs as bipartite graphs [28, 9], binding hyper-graphs may be depicted pictorially as in Figure 1:

**Figure 1:**
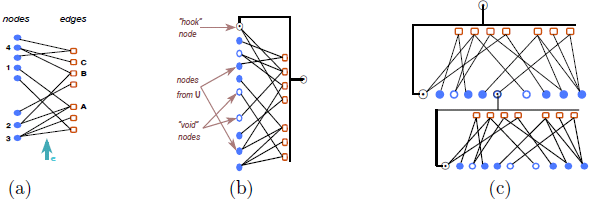
Hyper-graphs as Bipartite Graphs (a), Extended Hyper-graphs (b), and the Binding Prototype (c)

To complete the construction observe that, while there still are unbound meta-variables as nodes of any hypergraph in the assemblage obtained by binding extended hyper-graphs one to another, it is possible to indefinitely continue to bind other hyper-graphs to them. These assemblages are the whole-part graphs (*wp*-graph) and are formalized by the following recursive definition:

#### Definition 2.2

*An object γ is a* wp-*graph, that is, γ* ∈ **Γ** *if and only if:*

1. 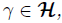
2. 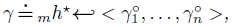
3. *nothing else.*

*Where *γ*° means that its “upmost” hyper-graph 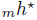 has a* ⊙ *meta-element as a node; that is*, 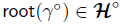. *And the symbol* ≐ *reads* is build as *or* is given by *and has a double interpretation: as a programming assignment during construction, then as a mathematical equality [13]*.

The fact that 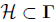 and a fixed point theorem for structures [19] warrant that **Γ** is well defined [13]. Note that Definition 2.2 does not make reference to sets. It can be proven, though, that **Γ** contains all its finite subsets. The possibility of indefinitely binding new *wp*-graph s to a chosen one means that these structures may grow forever and that adding details is unlimited.

### 2.3 Synexions

The elements of U can be a variety of things: letters, chemical atoms or molecules, forms, objects, concepts, ideas etc; the last two ones, as with any abstract object, if properly labeled by tokens, names or other signs. Hence, whole-part graphs from Definition 2.2 may represent organizations which (logical) atoms may be concrete, abstract or imaginary. As examples we have: words, paragraphs and text, molecules and macro-molecules, cultural entities, emotions, sketches, cognitive maps, pictures and anything we may name.

It is worth noting that the same object may be associated with several organizations, depending both on the objects’ intrinsic features to be highlighted and on the interests and purpose associated with their observation. For instance, the common organization of a text is grounded on sequences of letters, words, phrases, sentences, paragraphs, sections etc. This is the right organization for reading it. However, if we are interested in unveiling its key ideas and their associations a better organization would be as a cognitive map or a hypertext. Hyper-documents in the Web provide a variety of experiments in this direction, although not all them naturally conform to Definition 2.1.

Therefore, an object under study may be seen as different organizations, depending on the purpose and focus of our enquiries. Furthermore, there are things, like emotions, that may be both abstract and concrete: abstract while described by humans in words, or concrete whether we are considering the biochemical alterations and oscillations endowed by them. The organizations that may be associated to this sort of things, while taking them as abstract or concrete, are clearly profoundly different.

Biological (concrete) organizations, however, occupy volumes in space and our recognition of them is strongly molded by their spatial arrangements and their functional dependencies. Moreover, physical constrains may prevent, hamper or facilitate the existence of certain organizations in favour of others. Therefore, we need to provide means for representing organizations “embodied” in physical space. This will be achieved by synexions, or organized volumes, which for the purpose of the present text are simply defined as associations between elements of **Γ** and collections of elements in a “physical” space, satisfying a certain condition.

Thus, let 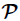 be a “physical” space, for instance, the Galilean space-time structure, a four dimensional affine space with a topology based on Galilean transformations [4]. And let 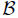 be the class of all associations between *γ* ∈ **Γ** and subsets of 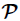, which will be denoted by 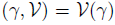, that preserve the hierarchy of *γ* in the collection of associated volumes. That is,

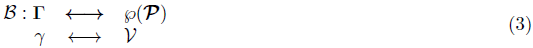

such that

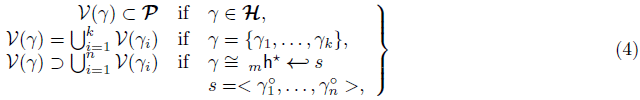

where *n ≤ m*.

Synexions are not sets in the usual sense. For one thing, subsets of 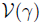 must also conform to conditions (4) and have parts of 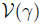 as their parts. Thus, we may have 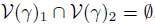, even though 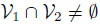 as subsets of 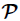. Moreover, many collections of volumes may be associated with just one *γ* resulting in several synexions, since conditions (4) do not univocally determine the associated volumes. A natural way of accomplishing many synexions out of a single *γ*, is to change the volumes associated with some parts of a *γ*, either by fattening or shrinking the volume itself or by changing its localization in space-time. This property of synexions allow them to represent movements and deformations of organized things more effectively because it is possible to impose a dynamic behaviour on 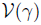 that change its associated volumes while preserving *γ*, their organizational identity. Cell-motion [2] is a good example, since organelles and cell-parts deform and move with the cell without destroying themselves. Synexions support the disentanglement of displacement and deformation from organizational changes.

### 2.4 Further Constructions

Synexions are not the only “concrete” organizations and the spaces **Γ** and 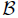 have many interesting properties and possibilities. This basic framework allows for development in many directions. Although a proper discussion about other features and more elaborated aspects of mathematical constructs concerning or associated with organizations, whole-part graphs, and synexions are outside the scope of this writing, two points should be noted here.

A process may be understood as a collection of *state* enchainments that are interpreted as entailments of natural events, that cohere enchainments and temporal order when instantiated in the physical space [16]. Since states in life phenomena are always organizations, processes are naturally casted as collection of enchainments in **Γ** with a condition about their immersion in the time dimension. Processes are thus organizations immediately representable in **Γ** and in 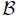. So, we may also consider organizations of processes. This is of great relevance while considering biological phenomena since chemical processes (as entailments of chemical reactions and substrates) are processes in the sense just described. Organizations of (bio-)chemical processes arise by considering two chemical processes to be associated whenever they exchange substrates and considering them as a whole whenever they present homeostasis and have a distinctive functional character.

Besides this, organizations modeled by *wp*-graphs have a natural measure of complexity grounded on counting and weighting possible and existing associations and counting the hierarchical levels induced by the recursion steps. This complexity measure is such that: the more associations there are in a *wp*-graph, the higher the complexity is. Moreover, the more detailed an organization is or the bigger the number of its parts is the higher its complexity is, because deeper the recursion steps go. Therefore, since the same natural object or entity may be associated with distinct organizations, it is a fortiori possible that they have different complexities. This means that complexity is a concept intrinsically more associated with the description and representation of a phenomenon in terms of organizations rather than with the phenomenon itself [14].

This introduction to whole-part graphs is rather terse, lacking important details. More formal descriptions of these constructs may be found in [13, 15] and forthcoming work.

## 3 Information

Organizations may convey information. This is clear whenever we consider texts or pictures as organizations and if their information content is conveyed to humans. Moreover, information-driven interactions are a distinctive feature of all biological systems, what is ingeniously discussed in [24]. However, existing information concepts may only be used to describe information-driven interactions in a very ad hoc basis and under constrains that keep important phenomena out of the description. A concept that may be used at all biological scales in an integrated manner and cover all relevant aspects of biological development is needed. Information conveyed by organizations fulfill this requirement, since it is tailored to biological interactions and the information conveyed is determined by organizational changes.

Information in the sense to be presented below is not a measure or even numeric. It goes towards the etymological source of the term information: *in-formare*, or “to form” inside. It is a concept that, nevertheless, can recast the usual numeric measures of information in special settings. In the following, we *assume* that any biological entity may be represented as a synexion, for a properly chosen atom set U.

### 3.1 Perceptions

The concept of information to be presented is grounded on an ontological hypothesis. Namely, that all biological entities “perceive”. The purpose of this section is to clarify the use of this term since it is used here with a rather specific meaning.

Perceptions are strongly intertwined with signals. Biological entities generally have a “skin” that delimits its boundaries. A skin is a physical region that divides the world in two regions: *inside* and *outside* the entity and is part of its organization. The “outside” region of an entity, together with anything contained in this region and that may interact with the entity itself, is its *environment*.

#### Definition 3.1

*A **signal** is any perturbation of the environment’s state, constituents, or attributes that is concentrated in a time interval and region of space and propagates, eventually encountering a biological entity*.

Encounters have the usual meaning of being at the same point (neighborhood) in space at the same moment. Due to space-time continuity, whenever a signal encounters a biological entity it clearly reaches its “skin” first. Perceptions are effects that a signal has upon a biological organization it encounters.

#### Definition 3.2

***Perception** is a two stage process: it has an imprint moment and a recall moment:*

▸ *Imprint* *Any signal **σ** encountering the sensory apparatus (in the skin) of a biological entity* 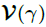 *at time t*_*σ*_ *and transmitted into it provokes a (localized) change in its organization*.
▸ *Recall* *Moreover, if another signal **σ**′ encounters the same biological entity at time t ≥ t_σ_ and tends to provoke the same change in its organization, it is recognized by synexion* 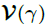 *as being the same perturbation as **σ***.

Hence, *perception* is an action rather than an entity, organization or fact and results in imprints (see Figure 2). Imprints may decay over a short time, remain for longer periods or become part of the organization. The signals ***σ*** and ***σ**′* need not be exactly the same perturbations of the environment when we take all their attributes into account. With respect to perception process of a synexion 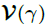, however, they will be the “same” whenever they associate to the same imprint. This is dependent on the complexity of 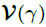 and those of signals in the imprint class associated with it. This sharpens the idea of similarity of signals.

**Figure 2:**
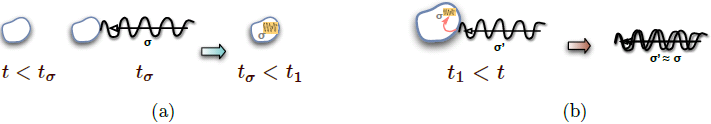
Perception: imprint (a), recall (b)

Mistaking strictly different signals related to a sole imprint as the same signal is part of the perception process. Therefore, we say that a long lasting imprint is a model for ***σ*** and its similarity class.

### 3.2 Information and interpretation

Imprints are changes in the organization of a synexion 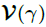. Imprints that change the behaviour of a biological entity will be considered to be in-formation. The definition below employ observers, that are biological entities themselves. The role of observers, notwithstanding, is primarily to acknowledge that something has happened. Since we will not be considering observation as an active/passive procedure, there is no point to discuss interference by observers. Observers are nevertheless needed whenever we want to discuss changes in any domain of discourse. Their role will be greatly clarified by making our understanding of what is information more precise.

The definition of information to be presented will rely on the following ontological hypothesis about living phenomena and entities:

#### Hypothesis 3.1

**A:** *Any biological entity or process may be represented as a **synexion***.

**B:** *All biological, or life related, entities or processes **perceive***.

**C:** *Perceptions are unique for a given biological entity or process that remains unchanged (same signal, same imprint);*

**D:** *All existing biological organizations emit a collection of signals that uniquely characterize them*.

Strictly speaking, hypothesis C and D are not necessary for the *definition* of information. They are though relevant while considering information as the kernel of biological interactions, for rendering information as a usable concept, and for clearly understanding its biological consequences.

It is worth to note that the representation of biological entities and processes as synexions is not unique or enforced in any manner. The same biological entity may be represented by synexions 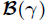 and 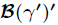 reflecting different levels of detail or different perspectives of study. Albeit just one representation is enough to admit discussing about information and the interpretation of signals. In consequence, studies involving information must carefully observe the representation chosen, because the imprint by a signal ***σ*** in 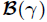 may be different from its imprint in 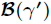 or 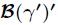. The similarity classes of ***σ*** may be even distinct for each of these representations. Our ability to understand information-driven interactions will thus strongly depend on how trustworthy the organizational changes due to perception are represented. This relates to the level of detail the synexion represents the inspected organization and a compromise between complexity and reliability will always exist. This is not strange to science but is made explicit in this setting.

Second, it is well accepted nowadays that cells perceive [21]. There is a clear association between environment conditions and activation-deactivation biochemical switches in the cell nucleus, maintained by the signaling cell system [3, 27]. The hypothesis 3.1 (B), however, goes beyond that towards the sub-cellular scale and above to non-organismic entities and collectivities scale. The present framework allows for treating information other than that processed by neural systems or DNA transcription and replication, that are the only forms of biological information generally considered till recently [24].

Before advancing further, is worth making the following observations:

- A signal ***σ*** reaching two different biological entities ***a*** and ***b***, represented by ***B***^**1**^(*γ*_1_)_***a***_ and ***B***^**2**^(*γ*_**2**_)_***b***_ may provoke different imprints in their organizations, even if *B*^1^, *B*^1^ and *γ*_1_, *γ*_2_ are similar. However, if the collection of similar signals is the same for both ***a*** and ***b***, that is, if any signal leading to the imprint caused by ***σ*** in *B*^1^(*γ*_1_)_*a*_ will also lead to the imprint caused by ***σ*** in *B*^2^(*γ*_2_)_***b***_, the perception should be considered same;
- A traveling molecule is a signal, because it is a variation in density, mass and other factors concentrated in time and space;
- Pressure and concentration variations, being caused by more diffuse and collective perturbations of environment attributes, may not always be taken as a signal. This suggests that environmental variations depend on scale sensitivities and the complexity of the signal and the perceiver to be considered as signals. Signal and perception are thus relative concepts;
- Encountering is always due to relative movements. Either the signal propagates or the organism is moving and reaches a perturbation in the environment fixed in space (like obstacles). What is important is that signal and organism approach each other in space and time for an encounter to occour.

Besides signals and perception, special biological entities will be required to specify information. One shall be termed an interpreter, the others will be observers.

#### Definition 3.3

*Given a signal **σ** and at least two biological entities* 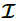 *and* 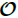, *σ will be termed an Information if the following things occour conjointly:*

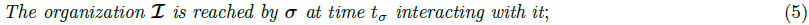

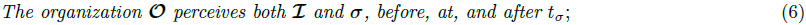

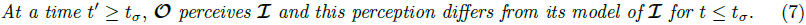

*That is, if* 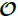 *perceives changes in* 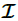*, after its encounter or interaction with σ*.

In this case, 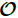 says that ***σ*** is an information for 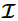 and that 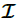 interpreted ***σ***. 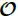 is just the observer that happened to acknowledge the interpretation. The observer 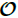 is not needed otherwise and 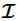 may be the observer itself if it is complex enough to be able to perceive its own signals or perceptions and maintain a model of itself.

Figure 3 pictures this information-interpretation definition, highlighting several observers (named 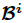 in the figure), models and the dependence in time.

**Figure 3:**
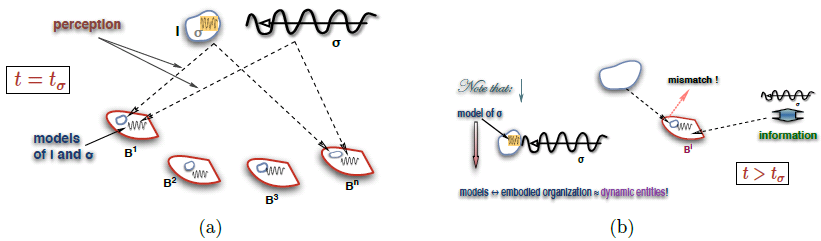
Information: Observing a signal perception (a); recognizing ***σ*** as information (b)

## 4 Space, Time and Information

This definition of information required the immersion of organizations in space-time, explicitly using spacetime events and space-time models in its definition. In the following, arguments supporting this dependence of information on space and time are presented at several scales and domains.

### 4.1 Cognitive Domain

Talking about issues of cognition and models starts with humans. This brings our analysis to a subject closer to our usual sense of information. Cognition, seen as acquisition of information and perceptions in the sense of Definitions 3.3 and 3.2, is not restricted to humans. In multi-cellular organisms, a part of their organization (the nervous system) is specialized as a signal processing system. It is dedicated as well to the treatment of the imprints resulting from perception and any response or reaction to them. Complex signals, coming out of complex organizations, are processed by the sensorial-signal systems and imprints are mostly associated to the brain and nervous system, although not restricted to them. In cells, an elaborated and complex network of reactions centered on DNA and nucleotides adapt and respond to signals [3, 11]. The examples below refer nevertheless to human cognition.

Signals are apparently affected by the organization of their sources. The organization of sources seems to be reflected in the imprints associated signals emanating from them, at least partially. In the following discussion, association of parts is mainly given by topological proximity (neighborhood), while the whole-part hierarchy by encapsulation of groups into unities.

In texts, that are sequential objects, the vicinity is given by collaterality, grouping letters into words. Words side by side from phrases. Phrases side by side, with punctuation marks interspersed, form sentences, and so on. However, parenthesis and footnotes relief a bit text dependence on sequential vicinity, by enclosing their contents into units. Technical texts have other forms of encapsulating and naming text unities that may be referred latter in the text, thus indirectly re-appearing at several points of the text.

The same mechanism appears in figures and pictures, although their inherent two or three dimensions (2D or 3D) make the idea of neighborhood much richer from a practical point of view. To exemplify how neighboring associate things, let’s consider two simple geometrical objects, here taken as wholes and logical atoms in U: a circle (◯) and a line segment (∣). The circle can be rotated at will without appearing to be modified but the line segment will present different inclinations if rotated. Bringing the circle and line segment together (see Figure 4) in different manners will reveal important characteristics of information.

**Figure 4:**
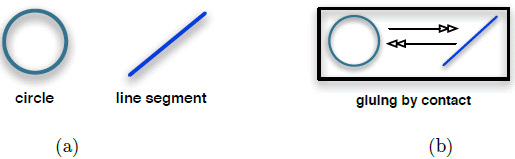
Geometrical objects (a); joining them by contact (b)

Note that the line segment may remain tangent to the circle after joining or may transect it and that the circumference of the circle may touch or cross the line segment at different distances form its center. These different possibilities of contact will result in different tokens, symbols and signs that may convey different meanings, if they convey information at all. Each circle-glued-with-segment forms now a visual unit (Figure 5), a whole in the sense of both **Γ** and 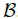.

**Figure 5:**
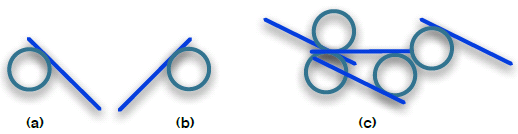
Circle-line-segment visual units

Not all wholes convey information, but there are cases where a whole may not convey information due to the manner it relates to its environment. For instance, it may be difficult to recognize anything or associate a meaning to the circle-segment wholes as they appear in Figure 5. There is no easy clue in the visual units of images (a) and (b) of Figure 5 or in those in the the heap of the image in Figure 5(c) that help us recognizing them. What, by definition 3.2, means associating these units (signals) to a previously imprinted (known) signal.

If we consider the images in Figure 5 (a) and (b) together as one image, it becomes possible to vaguely identify this new whole as pair of (angry) eyes… if one has seen *lots* of animations. But what if we straight these units and line them up like in Figure 6? Don’t they become immediately recognizable as letters {p,q,d,b}? Observe how implicit subliminal visual references to the borders of the paper, that provide a sense of verticality, and from one whole to the other enforce the identification of each one as a letter. It is easily observable, in logos or outdoors and other advertisement material, that a group of letters like a syllable can be rotated to any degree and be severely distorted remaining recognizable.

**Figure 6:**
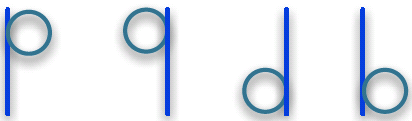
Letters

It is important to note that relative distances between letters retain strong topological internal clues that enforce groups of letters to act as units or wholes, reducing the relevance of clues related to the environment. Similar phenomena occour in time when we consider sounds and music. Groups of sounds or musical notes are more stable to our senses then scattered individuals sounds or notes. It is more difficult to make known music or meaningful spoken words unrecognizable by distorting them than individual sounds or new sound sequences.

Other visual units can be formed out a circle and a line segment, as shown in Figure 7. Some will be easily recognized like those of Figure 7 (a), (b) and (c); while others like (d) will not, even if due to very small relative displacements of one part in relation to the other. Possible meanings for the resulting symbols are indicated by labels within the picture.

**Figure 7:**
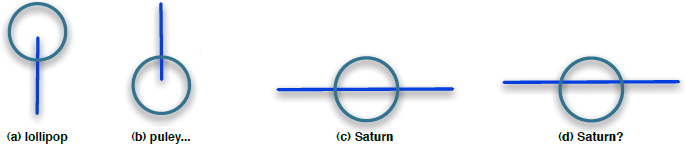
Other visual units out of a circle and line segment

From another stand, there are visual units that defy interpretation no matter what is done with them, like image (a) of Figure 8 in two dimensions. The image of Figure 8(b) may initially defy recognition. However, shaking or moving it a little bit, as indicated in Figure 8 (c) will make them recognizable. This recalls the importance of movement in recognition of many things and phenomena, particularly life. Synexions are space-time objects and tubes resulting from the displacement of any volume in time is a synexion. Hence, Definitions 3.2 and 3.3 contemplate cases where the recognition and interpretation of objects can only be made while they are moving.

**Figure 8:**
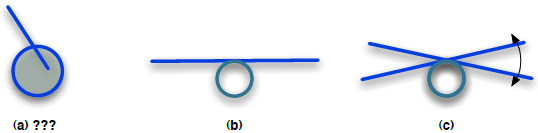
Visual units (wholes) difficult to recognize

Finally, visual assemblages and displacements that form wholes and have been recognized may be associated in infinitely distinct forms resulting in new units or wholes, as suggested in Figure 9. This process may be carried out indefinitely resulting in visual units composed of other visual units which recognition helps the recognition of the whole unit. Abstracting from top-down or bottom-up stands, the explicit recursion in Definition 2.2 and equation 4, that are inherited by Definitions 3.2 and 3.3, allows for the appearance of however complex organizations, signals, imprints and information.

**Figure 9:**
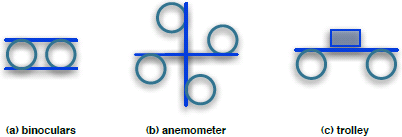
Visual units (wholes) difficult to recognize

### 4.2 Molecular Scale

The hierarchical structure of proteins is now established and they are considered to be formed by assemblages of secondary structures rather than sequences of amino-acids. The information about proteins and their structure is pretty large (see, for instance, [2] and the collection of articles at http://www.proteinspotlight.org for information and pictures). Proteins may be easily represented in **Γ**. Molecules are traditionally represented as graphs, or hyper-graphs. So, a whole protein can be depicted as a single graph, that would be awfully complicated. Given that the basic constituent parts of a protein are the amino-acids rather than atoms, a protein can be depicted as a sequence of the hyper-graphs representing them, and sequences of hyper-graphs belong to **Γ** [15]. Often, proteins are depicted in terms of their secondary structures. Considering the bounds that fold a sequence of amino-acids into secondary structures, we have a hyper-graph reflecting all these bindings. The node of which are hypergraphs, what by Definition 2.2 is in **Γ**. Therefore, any assemblage of secondary structures is also immediately modelled in **Γ**. Moreover, their representation in 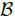 is also immediate, since the volumes and positions occupied by atoms and molecules in space, and their vibrations in space-time, give the required synexions 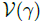 to represent proteins. It is also clear that any macro-molecule may have many different representations *γ* ∈ **Γ**, depending on which of its constituent parts are being considered as wholes. Consequently, macro-molecules have also multiple representations as 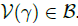.

At any point in time, a protein may be *activated* or *deactivated* by a chemical or other environmental signal reaching it. Activation and deactivation are due to changes in their organization either in **Γ**, if new chemical bonds are formed or small molecules become attached to the protein, or just in 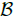, due to a re-arrangement in the tertiary structure caused by changes in the relative positions of the protein constituent parts. These organizational changes modify their “function”, that is the manner a protein chemically behaves and reacts to external stimuli. Therefore, a perturbation in the environment, be it a traveling molecule or variations in distribution of energy or mass concentrations, may cause an alteration in the protein organization that changes its behaviour. Under Definitions 3.2 and 3.3, the (chemical) perturbation is a signal that provokes a re-organization of the protein changing its chemical behaviour. Therefore, the perturbation is an information for the protein.

An example of organization alteration due sole to changes in space and not in **Γ**, is the pigmentation protein chameleonine [10]. Changes in environmental conditions, in this case temperature variations, provoke a change in the color of the tissue where it resides. Now, temperature variations can only change the energy of its components that reside in the vibrations of the protein’s atoms. Chameleonine responds to this with a deformation in its spatial configuration due to the consequent structural stresses. As a result of this new (spatial) organization, it changes behaviour reflecting a different light frequency. The signal in this case is the temperature variation that, at the molecular scale, means a change the mean number of molecules hitting it per time slice. This is an example that conspicuously show the need to include space and time in the conceptualization of information.

Another example of spatial dependence of perception and information is chiral isomers. This is a rich subject that will not be further discussed here due to size constrains. However, a few observations will illustrate important aspects of perception and information. Chiral isomers commonly respond equally to simple stimuli like temperature or to simple substrates in chemical reactions. However, depending on the complexity of molecules involved, on the environment where they are immersed or on the complexity of the signals reaching them, the effect in the organization of wholes of which they are parts may be dramatically distinct, provoking deforming developmental diseases in humans, like the infamous thalidomide, particularly when chiral isomers are the signals [1, 17]. Remind that life phenomena is strongly dependent on essentially one of the chiral isomers.

From another stand, many biochemical organizations and structures in cells are not static. They are grounded on interacting homeostatic chemical processes, particularly when cycling. Hence, biological phenomena really build on organization of processes. A now canonic example of this character of theirs are bacteria molecular motors. There are pictures of these ingenious engines available. However, these “structures” do not appear in state snapshots at a fixed moment in time. They arise from the superimposition of several snapshots [6, Figure 2] at distinct times meaning that a flux of chemical substrates cooperate to give existence to these “motors”. In term of organizations and synexions, notwithstanding, they are naturally represented and may also be observers and interpreters.

### 4.3 Cellular Scale

At the cellular scale, organizations are more conspicuous in eukaryotes than in prokaryotes, in spite of the complex organization of biochemical processes existing in the latter. The preferred reference to organizations in eukaryotes in the sequel is just a matter of explanatory simplification.

Looking into a cell structure, there are organelles, special “tissues” like the endoplasmic reticulum and the vacuoles’ membranes, as well as a plethora of sustained biochemical processes that serve a variety of purposes in the cell. Among them we find transportation systems that using vacuoles protect substrates from reacting with chemicals existing in the cell before they reach a certain place or organelle in the cell. Signaling systems, extending from the external to the nucleus membranes, are responsible to transmit changes in the state of chemical switches in cell membranes, that act as sensors, to the nucleus. There, the nucleotide-DNA system retain the switches conditions in the form of inhibition-activation patterns, or imprints (Definition 3.2).

This is mainly a traveling wave system that changes the switching status of a collection of molecules that surrounds the DNA and maintain some portions of the genes active while inhibiting others. Besides messenger RNA, that transport information, there are also signaling pathways from the nucleus to the cytoplasm that activate or deactivate protein synthesis, that are determined by the inhibition-enhancing patterns of DNA sites. Therefore, it is not DNA that really controls cell activities but a complex formed by DNA and nucleotides that “switch” genes on and off. The messenger and transcriptions systems transmit these patterns of free DNA to appropriate sites in the cell. Also in this case, environmental perturbations that change the status of membrane sensors are signals as in Definition 3.1, while the DNA-nucleotide complex act both as memory (the organization that concentrate imprints) and a transducer that may affect cell behaviour, that is, interprets (Definition 3.3).

To see biotic-interactions as exchange of information, we need to understand the effects of differences in time propagation between two wholes. In prokaryotes, transportation is due mainly to diffusion in heterogeneous media, that takes little time to bring some molecule from one extreme of the cell to another, due to their small volume. Eukaryotes are about 10 times as lager as prokaryotes. To get equivalent transfer and reaction rates and characteristic times in larger cells, there must be some facility to “direct” and accelerate diffusion. This “facility” is organization. More precisely, dynamical organizations.

### 4.4 Physiology and Behaviour

Comments in this section refer to multi-cellular organisms. Scales up, the organization of multi-cellular organisms pretty ressembles that of eukariotes and prokariotes from the right perspective. Following J. Miller, apud Peacoke [20, 23], living systems are composed of nineteen systems, independently of its scale. For instance, there is the nervous (electrical signaling) system, the immune (repair) system, the digestive system, the motor system, the boundary (environmental interface) system, the memory and learning systems. In spite of being analogous, they may however appear in completely distinct forms from one scale to another.

Par contre, there are systems in larger organisms that are not as easily paired among scales as the former, like the endocrine system (chemical signaling?) or the boundary systems in ecosystems and societies. Moreover, the categorization in nineteen systems is somewhat arbitrary, since some may be considered as one and others decomposed. Also, focusing on the nervous and endocrine systems, we see that two signaling systems with remarkably different characteristic propagation times co-exist, one based on electro-chemical wave propagation and the other on diffusion and transport. Moreover, phenomena occurring within the limits of these systems is much richer and complex for multi-cellular organisms than for cells and, by extrapolation, scales up.

Instead of systems, multi-cellular organisms and more complex organisms are better represented as a collection of superimposed organizations symbiotically cooperating by exchanging matter, energy and information, particularly the latter. As in the case of systems, a categorization of which these organizations should be will always be arbitrary to a certain extent and finding biologically sound heuristics for this will greatly enhance our knowledge about such systems. Clearly the macroscopic size of multi-cellular and more complex organisms impose a strong dependence on space and time for any exchange within and among the organizations composing a living system. Therefore, rendering the categorized organizations compatible in terms of characteristic times, volumes and frequencies is a mandatory guideline while discovering these organizations.

When we acknowledge the exchange of information as a distinguishing characteristic of living matter, the necessity of this compatibility greatly promotes the use of space and time in the definition of information.

### 4.5 Cultural Domain

Contrary to knowledge that may be individual, culture is a collective phenomena. Notwithstanding, human purposeful collectivities (like enterprises, firms, industries, households, schools, communities etc), collective phenomena (like person-to-person contacts that propagate diseases, gossips, information and matter; or like cultural centers and cultural networks attracting and educating people) and human creations (like science, music, art and literature) are all founded on organizations and information as pictured in sections 2 and 3.

Purposeful human collectivities are formed by a collection of people that associate with each other with a (pre-)determined objective. Some organizations, like firms, industries, enterprises and (big) households, spontaneously or not organize themselves by having a hierarchy where smaller groups report to and/or are commanded by others. Organizations as such are the basic action units in a cultural domain and have been extensively studied since long [29]. Most of them comply at least approximatively if not strictly to Definition 2.1.

Knowledge may be represented by classes of signals and their imprints organized as a consequence of associations and abstractions which are formed out of multiple interpretations and experiences, in the lines suggested by the discussion of subsection 4.1. Culture also may be viewed as an organization (in the sense of Definition 2.1) composed of concepts, ideas, taboos, individual knowledge, the imprints of a man or population etc. For instance, in a piece of music, art or literature there are often references, usually implicit, to another work in the same class or even in another class. There is music inspired by literature and folkloric wisdom, literature that refer to lyrics and so on. Literature is itself a web of veiled references to other literature.

In science, referencing is made explicitly; and this differentiates it from the other cultural expressions. Books, although sequentially arranged, may be as tortuous as any folding molecule due to backward and forward cross-referencing [25, Note to the reader]. In either science, literature or music, one can only appreciate the beauty and depth of a work if one has a good acquaintance with a large number of other work, at least in the same cultural class. Even within the explicit referential system of science, one has to know and understand the referred material to properly perceive and understand the work that referred to them.

Culture, however, differs from individual knowledge in two important ways. Instead of residing in the memory of a person, it is registered in books, writings, unspoken cultural premisses, “learning by seeing” or by “experiencing” and many other extra-organic media and conveyors. Furthermore, the interpretation of this collection of signals and imprints is not made by an individual in particular but, instead, by the whole community that produces and retains the culture. This means that many of the writings and extra-organic, non-individual, media contain ideas and discussion about elements of the culture itself, being self-reflexive and self-referential to a much higher extent than knowledge.

The intrinsic nature of science is not much different than that of culture. Its findings (atoms in U) are also organized in the manner of Definition 2.1 and its interpretation is a collective enterprise. The distinctions reside primarily on rules, methods and underlying paradigms about how to collectively develop scientific knowledge and the election of observation as a resolver of interpretation and argument conflicts. The language of science, even when it doesn’t use mathematics, is developed in such a manner that differences in interpretation are minimized, arguing and interpretation procedures are standardized, and the rules governing observation, interpretation and argumentation accessible to everyone. In this sense, the collective doing of science is more self-conscious than that of culture. Culture is produced more instinctively and intuitively than science, where there is a more definite and explicit collective purpose to be followed.

## 5 Conclusions

This is a quick description of a fairly sophisticated mathematical framework that is able to represent (biological) organization and information as conveyed by organizations, followed by arguments justifying the immersion of organizations in space and time to support the procedure of defining information. The information concept presented in this paper is non-quantitative and is considered to be biological solely in terms of the perception hypothesis. This is an ontological assumption (supposedly) fulfilled by any biological entity, even at sub-cellular levels. Nowhere else does biological aspects intervene. Hence, it may be used in situations where an interpretation of perception, in the specific sense described, is available. A detailed account of this mathematical framework is in preparation.

It is worth noting a few points about these definitions. First, time and space are crucial for defining information not organization. Time and space differentiation, variation, and scale integration are at the very kernel of many biological information-driven interactions. Therefore, there may be organizations that convey information to nobody (and this may remain true even if they encompass space and time). Furthermore, there is room in the framework evolving from the definitions above to handle and study attributes and characteristics taken to be exclusive of life phenomena, and to tackle several current conflicts and misunderstandings about the use of information to describe and explain life phenomena.

Second, an interpreter may be himself the observer. This requires that its complexity be high enough to house a model of itself and of the signals it receives. The alluded complexity is intrinsic to each organization and relates to the number of its parts, connections, and hierarchy levels [13].

Third, the parts of observers that model the interpreter, the signals, and interactions resulting in interpretations are also synexions, i.e., organized space-time volumes within the observer 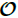. Besides this, observers record signals, encounters, interactions and interpreters in space **and** time. Hence, the models observers keep of signals and interpreters have projections along time, encompassing “expectations” about their behaviour, at least towards an immediate future. This allows considering the treatment of purpose and teleologic actions like escaping from approaching objects in collision paths and *anticipation* [25, 22].

To complete a basic framework for the description of life, though, a path towards a “dynamics” of organizations must be constructed, that is, a dynamics able to describe enchainments of *changes* in organizations *γ* rather in the volumes of 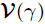, and infer thereupon. A step in this direction is to consider, by analogy to molecular and physical dynamics, some form of fluctuation of organizations. Although a more detailed account of the ways this can be represented in this framework is outside the scope of this text, requiring a much deeper immersion in the underlying mathematical ideas just presented, it should be noted that, with respect to information and information acquisition, either from a bio-chemical [18], communal [8] or other stand, organization fluctuations may hamper or facilitate information acquisition and learning processes through distinct effects, depending on being syntonic or distonic to various degrees. It is thus, in this sense, related to the most basic (physico-chemical) expression of emotions.

## Acknowledgements

These ideas and framework have been developed and polished during a few decades. During this time, I had the pleasure to be assisted by a great number of colleagues and people in the long standing process of consolidating these ideas, through discussions or support of variegated kinds. The list presented in the sequel is certainly incomplete, due to the duration of this work. My heartfelt apologies to those not in the list. I would like to express my deepest gratitude to the whole group of Mathematical Modeling of Knowledge Diffusion, headquartered at LNCC and the Federal University of Bahia, to the Faculty of Life Sciences of the University of Manchester, as well as, to J.-Y. Béziau, A.C. Gadelha Vieira, M. Grinfeld, S. Hubbard, F. Lopes, S. McKee, A.C. Olinto, N. Papavero, M.M. Peixoto, A.A. Pinto, A. Prokop, D.L. Robertson, M. Trindade dos Santos, J.-M. Schwartz, I.W. Stewart, S. Webb, and my present and former students, many of whom are current collaborators in research projects where these ideas are being applied. I would also like to acknowledge the financial support of the following agencies and programs: CAPES, PCI/LNCC, and FAPERJ (Grant 101.261/2014).

## References

[1] Chunzhi Ai, Yan Li, Yonghua Wang, Yadong Chen, and Ling Yang. Insight into the effects of chiral isomers quinidine and quinine on cyp2d6 inhibition. Bioorganic & Medicinal Chemistry Letters, 19(3):803–806, Jan 2009.

[2] Bruce Alberts, Alexander Johnson, Julian Lewis, Martin Raff, Keith Roberts, and Peter Walter. Molecular Biology of the Cell. Garland Science, New York, NY, 4th edition, 2002.

[3] Uri Alon. An Introduction to Systems Biology: Design Principles of Biological Circuits. Mathematical and Computational Biology. Chapman & Hall/CRC, London, 2007.

[4] Vladimir Igorevich Arnol’d. Mathematical Methods of Classical Mechanics, volume 60 of Graduate Texts in Mathematics. Springer-Verlag, Berlin, 1978.

[5] John Barwise. Admissible Sets and Structures. Springer-Verlag, Berlin, 1975.

[6] H Berg. Motile behavior of bacteria. Physics Today, 53(1):24–29, Jan 2000.

[7] Claude Berge. Graphs and Hypergraphs. North-Holland, Amsterdam, 1973.

[8] Hernane Borges de Barros Pereira, Mario Cezar Freitas, and Renelson Ribeiro Sampaio. Fluxos de informações e conhecimentos para inovações no arranjo produtivo local de confecções em salvador/ba. DataGramaZero, 8(4), Agosto 2007.

[9] Marcelo Trindade dos Santos and Maurício Vieira Kritz. On the hierarchical organization of metabolic networks: an underlying mathematical model. In Rubem Mondaini, editor, BIOMAT 2005 - International Symposium on Mathematical and Computational Biology. Selected Contributed Papers, Rio de Janeiro, 2006. e-papers Editora.

[10] Vivienne Baille Gerritsen. Moody wallpaper, April 2003.

[11] Franklin M. Harold. The Way of the Cell: Molecules, Organisms and the Order of Life. Oxford University Press, Oxford, 2001.

[12] Franklin M. Harold. Molecules into Cells: Specifying Spatial Architecture. Microbiology and Molecular Biology Reviews, 69(4):544–564, December 2005.

[13] Maurício Vieira Kritz. On biology and information. P& D Report 25/91, LNCC/MCT, Petrópolis, December 1991.

[14] Maurício Vieira Kritz. Creating bio-mathematical worlds. P&D Report 29/95, LNCC/MCT, Petrópolis, November 1995. (work presented at 13th European Meeting on Cybernetics and Systems Research, Vienna, Austria, April 9-12, 1996).

[15] Maurício Vieira Kritz. On relations between i-graphs and data structures. Logique et Analyse, 39(153–154):153–164, 1996.

[16] Maurício Vieira Kritz. Biological organizations. In Rubem Mondaini, editor, Proceedings of the IV Brazilian Symposium on Mathematical and Computational Biology — BIOMAT IV, Rio de Janeiro, 2005. e-papers Editora.

[17] H.Z Levinson and K Mori. The pheromone activity of chiral isomers of trogodermal for male khapra beetles. Naturwissenschaften, 67:148–149, Apr 1980.

[18] D Q M Madureira, L A V Carvalho, and E Cheniaux. Attentional Focus Modulated by Mesothalamic Dopamine: Consequences in Parkinson’s Disease and Attention Deficit Hyperactivity Disorder. Cognitive Computation, 2(1):31–49, November 2010.

[19] Zohar Manna. The Mathematical Theory of Computation. McGraw-Hill Co., N. York, NY, 1974.

[20] J. G. Miller. Living Systems. McGraw-Hill Book Co., Inc., N. York, NY, 1978.

[21] Amir Mitchell, Gal H Romano, Bella Groisman, Avihu Yona, Erez Dekel, Martin Kupiec, Orna Dahan, and Yitzhak Pilpel. Adaptive prediction of environmental changes by microorganisms. Nature, 460(7252):220–224, July 2009.

[22] Mihai Nadin. Anticipation and dynamics: Rosen’s anticipation in the perspective of time. International Journal of General Systems, 39(1):3–33, Feb 2010.

[23] Arthur Robert Peacocke. An Introduction to the Physical Chemistry of Biological Organization. Clarendon Press, Oxford, 1983.

[24] Juan G. Roederer. Information and its Role in Nature. The Frontiers Collection. Springer Verlag, Berlin, 2005.

[25] Robert Rosen. Life Itself: A Comprehesive Inquiry into the Nature, Origin, and Fabrication of Life. Complexity in Ecological Systems Series. Columbia University Press, New York, NY, 1991.

[26] Marta Sales-Pardo, Roger Guimera, Andre A Moreira, and Luis A Nunes Amaral. Extracting the hierarchical organization of complex systems. Proceedings Of The National Academy Of Sciences Of The United States Of America, 104:15224–15229, 2007.

[27] Tapesh Santra, Walter Kolch, and Boris N Kholodenko. Navigating the multilayered organization of eukaryotic signaling: a new trend in data integration. PLoS Computational Biology, 10(2):e1003385, February 2014.

[28] Gunther Schmidt and Thomas Ströhlein. Relations and Graphs: Discrete Mathematics for Computer Scientists. EACTS Monagrphs on Theoretical Computer Science. Springer-Verlag, Berlin, 1993.

[29] Herbert Alexander Simon. The Sciences of the Artificial. The MIT Press, Cambridge, MA, 3rd edition, 1996.

